# Preterm birth accelerates the maturation of spontaneous and resting activity in the visual cortex

**DOI:** 10.1101/2023.01.20.524993

**Authors:** Isabelle F. Witteveen, Emily McCoy, Troy D. Holsworth, Catherine Z. Shen, Winnie Chang, Madelyn G. Nance, Allison R. Belkowitz, Avery Dougald, Meghan H. Puglia, Adema Ribic

**Affiliations:** Department of Psychology, College and Graduate School of Arts and Sciences, University of Virginia, Charlottesville, VA 22904; Program in Fundamental Neuroscience, University of Virginia, Charlottesville, VA 22903; Department of Neurology, School of Medicine, University of Virginia, Charlottesville, VA 22903

**Keywords:** Preterm, Brain, Visual, Inhibition, EEG, Neuron, Activity, Aperiodic

## Abstract

Prematurity is among the leading risks for poor neurocognitive outcomes. The brains of preterm infants show alterations in structure and electrical activity, but the underlying circuit mechanisms are unclear. To address this, we performed a cross-species study of the electrophysiological activity in the visual cortices of prematurely born infants and mice. Using electroencephalography (EEG) in a sample of healthy preterm (N=29) and term (N=28) infants, we found that the maturation of the aperiodic EEG component was accelerated in the preterm cohort, with a significantly flatter 1/f slope when compared to the term infants. The flatter slope was a result of decreased spectral power in the theta and alpha bands and was correlated with the degree of prematurity. To determine the circuit and cellular changes that potentially mediate the changes in 1/f slope after preterm birth, we used *in vivo* electrophysiology in preterm mice and found that, similar to infants, preterm birth results in a flattened 1/f slope. We analyzed neuronal activity in the visual cortex of preterm mice (N=6 preterm and 9 term mice) and found suppressed spontaneous firing of neurons. Using immunohistochemistry, we further found an accelerated maturation of inhibitory circuits. In both preterm mice and infants, the functional maturation of the cortex was accelerated, underscoring birth as a critical checkpoint in cortical maturation. Our study points to a potential mechanism of preterm birth-related changes in resting neural activity, highlighting the utility of a cross-species approach in studying the neural circuit mechanisms of preterm birth-related neurodevelopmental conditions.

## Introduction

Preterm infants born <37 weeks’ gestation currently represent a tenth of the infant population, with rising prevalence. Risks for prematurity include multiple gestations and pregnancy complications, with health outcomes positively correlated with increasing age at birth (Blencowe et al., 2012, 2013; Barfield, 2018). As the brain is still developing during the third postnatal month, premature birth can interrupt the trajectory of multiple developmental processes in the brain, resulting in neurological conditions, developmental delays, and visual deficits (Limperopoulos et al., 2007; Ortinau and Neil, 2015; Leung et al., 2018; Mann et al., 2011). Neurocognitive and behavioral conditions are common in preterm children and adults, with a high prevalence of attention disorders and autism spectrum disorder (Limperopoulos et al., 2007; Strang-Karlsson et al., 2010; Treyvaud et al., 2013; Del Hoyo Soriano et al., 2020; Jaekel et al., 2013; Shuffrey et al., 2022; Crump et al., 2021; McGowan and Sheinkopf, 2021).

At birth, the brains of preterm infants are significantly smaller and often display persistent abnormalities in functional connectivity (Bouyssi-Kobar et al., 2018, 2016; Pandit et al., 2014; Hunt et al., 2019; Tokariev et al., 2019). Poor neurocognitive outcomes are, in some cases, associated with reduced connectivity in the brains of preterm infants and children (Pandit et al., 2014; Wehrle et al., 2018; Jakab et al., 2020; Ball et al., 2013; Kozhemiako et al., 2019). On the other hand, sensory cortices of preterm infants show increased connectivity when compared to *in utero* fetuses of the same gestational age, suggesting that preterm birth alters developmental trajectories in a circuit-specific manner (De Asis-Cruz et al., 2020). Given the increasing prevalence of preterm birth, identifying the changes in brain structure and function resulting from preterm birth and their mechanisms is vital for understanding the drivers of poor neurocognitive outcomes in the preterm population.

Electroencephalography (EEG) has been increasingly used in the clinical setting both for prognostic and diagnostic purposes (Holmes and Lombroso, 1993; Rivera et al., 2021; Watanabe et al., 1999; McCoy and Hahn, 2013). EEG is non-invasive and as little as 40 s of recording is sufficient to construct basic measures of neural activity, such as neural oscillations (periodic) and background (aperiodic) neural activity (O’Toole and Boylan, 2019). Both periodic and aperiodic components of the EEG power spectra undergo developmental changes, with spectral power decreasing in low frequency power bands and the slope of aperiodic component flattening with increasing age (Schaworonkow and Voytek, 2021; Gasser et al., 1988; Saby and Marshall, 2012). Both components have been used as a diagnostic and prognostic tool in the preterm population, with varying success (O’Toole and Boylan, 2019; Lloyd et al., 2021; Vanhatalo et al., 2002; Watanabe et al., 1999; Nishiyori et al., 2021; Shuffrey et al., 2022; Nordvik et al., 2022). Early postnatal EEG correlates with neurocognitive outcomes in childhood and the aperiodic component is associated with autism risk (Nordvik et al., 2022; Cainelli et al., 2021; Shuffrey et al., 2022). It is currently unknown how preterm birth affects EEG components in healthy preterm born infants when compared to corrected age-matched term infants.

Increasing use of animal models that replicate electrophysiological signatures of preterm birth-related brain injury, such as hypoxia, has aided in identifying the cellular and circuit changes after preterm birth (Zanelli et al., 2014; Burnsed et al., 2019; El-Hayek et al., 2011; Johnson et al., 2022). Very early preterm infants and hypoxic mice share common neural deficits, such as impaired development and integration of cortical interneurons (Stolp et al., 2019; Scheuer et al., 2021; Lacaille et al., 2019; Malik et al., 2013). Animal models of preterm birth itself are less commonly studied, likely due to low viability (Elovitz and Mrinalini, 2004; Dudley et al., 1996; McCarthy et al., 2018). Previous research has shown that preterm mice do not display impairments in tasks assaying anxiety-like and social behaviors, or in basal synaptic transmission in the hippocampus (Chiesa et al., 2019). However, impairments of visual processing and visuo-spatial attention are more common in preterm children (Leung et al., 2018; Burstein et al., 2021; Fazzi et al., 2012; Chokron et al., 2021), but it is currently unknown how prematurity affects the activity in the primary visual areas. An animal model that replicates the effects of prematurity on neural activity, especially in sensory areas, would be crucial for identifying the drivers of poor neurodevelopmental outcomes in preterm population.

In this study, we used a cross-species approach to determine the effects of premature birth on neural activity in infants and mice at a network level, and then at a circuit and cellular level in preterm born mice. We used EEG in healthy preterm infants and *in vivo* electrophysiology in preterm mice to identify changes in periodic and aperiodic neural activity associated with prematurity. We focused our study on the visual cortex for three reasons: 1) it is well characterized anatomically and functionally in both humans and mice (Kiorpes, 2016; Huttenlocher et al., 1982; Pinto et al., 2010; Haak et al., 2015; Norcia et al., 1987; Lunghi et al., 2011; Mitchell and Maurer, 2022; Shen and Colonnese, 2016; Colonnese et al., 2010; Antonini et al., 1999), 2) previous research suggests that the maturation of visual areas is accelerated after preterm birth (Schwindt et al., 2018; De Asis-Cruz et al., 2020; Jandó et al., 2012), and 3) the aforementioned incidence of visual impairments in preterm born children. We find that the aperiodic 1/f component slope is significantly flatter in both preterm infants and mice, confirming accelerated maturation of visual areas. Using preterm mice, we identify increased inhibition in the preterm brain as a potential mechanism of preterm birth-related changes in neural activity.

## Method

### Infants

Sixty-eight preterm infants recruited from the University of Virginia neonatal intensive care unit (NICU) and 75 term infants recruited from the greater Charlottesville area completed a resting-state EEG paradigm as part of a larger, ongoing study. Inclusion criteria for the preterm sample were birth prior to 37 weeks’ gestation, corrected age 0-4 months at testing, English-speaking legal guardian, no known uncorrectable severe auditory or visual deficit, and health condition deemed sufficiently stable for participation by the neonatology care team. Inclusion criteria for the term sample were birth after 37 weeks’ gestation, age 0-4 months at testing, English-speaking legal guardian, no known family history of neurodevelopmental disorder, and no known uncorrectable auditory or visual deficit. All data were collected at the onset of the critical period for the development of binocularity (after birth) (Fawcett et al., 2005), a visual function that is highly sensitive to altered perinatal experience (Freeman and Ohzawa, 1992; Fawcett et al., 2005; Jandó et al., 2012). The infant’s legal guardian provided written informed consent for a protocol approved by the University of Virginia (UVA) Institutional Review Board (HSR210330 or HSR19514, principal investigator: Puglia). Families were compensated $50 for their participation.

### EEG acquisition and preprocessing

Full scalp resting-state EEG was recorded while the infant rested in a caregiver’s arms for up to 7 min. Data were preprocessed with an automated preprocessing pipeline specifically validated and shown to generate reliable estimates for the computation of aperiodic signal in pediatric EEG data (APPLESED, Puglia et al., 2022). See Supplemental Methods for additional details. After preprocessing, 29 preterm and 28 term infants had sufficient data for subsequent analysis. See Table 1 for participant demographic and perinatal characteristics.

**Table 1.** Participant demographic and perinatal characteristics.

|  | **Preterm infants**<br>(n = 29) | **Term infants**<br>(n = 28) |
| --- | --- | --- |
| GA (weeks), mean ± SD | 31.4 ± 3.4 | 39.2 ± 0.9 |
| Extremely preterm (<28 weeks) | 6 (20.7%) | - |
| Very preterm (28-32 weeks) | 8 (27.6%) | - |
| Moderate preterm (32-34 weeks) | 4 (13.8%) | - |
| Late preterm (34-37 weeks) | 11 (37.9%) | - |
| Early term (37-39 weeks) | - | 6 (21.4%) |
| Full term (39-41 weeks) | - | 22 (78.6%) |
| Late term (>41 weeks) | - | - |
| PMA at test (weeks), mean ± SD | 39.0 ± 5.4 | 47.9 ± 4.6 |
| CA at test (weeks), mean ± SD | 7.5 ± 6.8 | 8.7 ± 4.4 |
| Sex |  |  |
| Female | 16 (55%) | 15 (54%) |
| Male | 13 (45%) | 13 (46%) |
| Race |  |  |
| White | 23 (79%) | 21 (75%) |
| Black | 5 (17%) | 1 (3.5%) |
| More than 1 race | 1 (3%) | 5 (18%) |
| Unknown race | - | 1 (3.5%) |
| Delivery method |  |  |
| Vaginal | 9 (31%) | 18 (71%) |
| Cesarean | 20 (69%) | 7 (25%) |
| Unknown | - | 1 (4%) |
| Birth weight (grams), mean ± SD | 1681 (664) | 3652 (817) |
| SGA | 6 (20%) | - |
| LGA | 3 (10%) | 5 (18%) |
| Obstetric complications |  |  |
| Third-trimester bleeding | 4 (14%) | - |
| Hypertensive disease of pregnancy | 11 (38%) | 1 (4%) |
| Rupture of membranes | 6 (21%) | 2 (7%) |
| Fetal growth restriction | 6 (21%) | - |
| Breech position | 3 (10%) | 3 (11%) |
| Maternal substance use | 2 (7%) | 1 (4%) |
| Respiratory issues | 23 (79%) | 12 (43%) |

| Other | 9 (31%) | 5 (18%) |
| --- | --- | --- |
| Antenatal steroid administration | 21 (75%) | - |
| Retinopathy of prematurity | 11 (38%) | - |
| APGAR at 5 min, median (min, max) | 7 (2, 8) | 8 (6, 9)* |

All data were extracted from hospital records when available (29 preterm; 24 term), or self-report (0 preterm; 3 term). Data for 1 term infant was not provided. Data are expressed as sample size unless otherwise stated. GA, gestational age; PMA, postmenstrual age; CA, chronological age; SGA, small for gestational age, defined as $< 10^{\text{th}}$ percentile; LGA, large for gestational age, defined as $>10^{\text{th}}$ percentile. Other obstetric complications include amniotic and placental complications, maternal fever/obesity/metabolic disease, fetal tachycardia, hypotonia, and nuchal cord. \*Apgar scores were not available for 4 term infants.

### Mice

Mice were maintained on C57BL/6 background (The Jackson Laboratory, Bar Harbor, ME) on standard 12:12 light:dark cycle, with food and water *ad libitum*. Animals from both sexes were used during the 4^th^ week after birth. Preterm mice were generated through timed breedings, where the day after the pairing was considered as gestational day (GD) 0. Once the pregnancy was confirmed (>1.5g increase in weight at GD 10), pregnant dams were habituated to handlers by daily handling. Mifepristone (MFP, Milipore Sigma, Burlington, MA) was dissolved in DMSO and 150 μg was injected subcutaneously on GD 17 (Figure 2A)(Dudley et al., 1996). Preterm mice were delivered on GD 18 (0.75-1 day early depending on the precise parturition time). Preterm mice have a significantly lower birth weight (Figure 2B), and increased mortality rates due to hypothermia, hypoxia, and inability to suckle (1-3 pups/litter). The cage with preterm mice was therefore supplemented with external heat and occasional oxygen to prevent hypothermia and hypoxia. Surviving pups are otherwise viable and display a catch-up growth typical of preterm infants (Figure 2C) (Altigani et al., 1989). Preterm mice open their eyes significantly earlier, further suggesting accelerated development of visual brain areas after preterm birth (Figure 2D). Control term mice were obtained from timed pregnant dams injected with DMSO only on GD 17. To match the infant cohort as best as possible, all mice were tested in their fourth week of postnatal development as that period represents the onset of the critical period for binocular maturation in mice (Wang et al., 2010). Animals were treated in accordance with the University of Virginia Institutional Animal Care and Use Committee guidelines.

### In vivo electrophysiology in mice

V1 recordings were performed on awake term (N=9) and preterm (N= 6) female and male mice, ages 21 to 28 days after birth, using a treadmill and blank screen as described in (Niell and Stryker, 2010). See Supplemental Methods for additional details.

### Immunohistochemistry and imaging

Brains from term and preterm mice (aged 35-40 days, N=3-6/group, as indicated in text and figure legends) were sectioned into 40 μm, stained, and mounted on glass slides. Images were acquired from 4-6 sections/mouse (minimum 20 images/mouse). Quantification was performed on background subtracted images using ImageJ. See Supplemental Methods for additional details.

### Quantification and statistical analysis

Statistical analyses were performed blind to condition in GraphPad Prism 9.0 (GraphPad Inc., La Jolla, USA) using nested t-test and one or two-way ANOVA with post-hoc comparisons (as indicated in text and figure legends), unless stated otherwise. Spectral analysis was performed with Spike2 on 60 s of data at 500 Hz for both infant and mouse datasets. Power was normalized to the power in high frequency band (150-250 Hz). For infant data, power between 58 and 62 Hz was excluded to eliminate contamination from line noise which was present in some samples. Power spectral density values were averaged across channels of interest (Figure 1A). 1/f slope was generated using linear regression from 2-25 Hz for infants and 2-70 Hz for mice after log-transform of frequency and power. All data are reported as mean ± SEM, where N represents number of animals and infants used, unless indicated otherwise. Target power for all sample sizes was 0.8 and alpha was set to 0.05.

**Figure 1.**
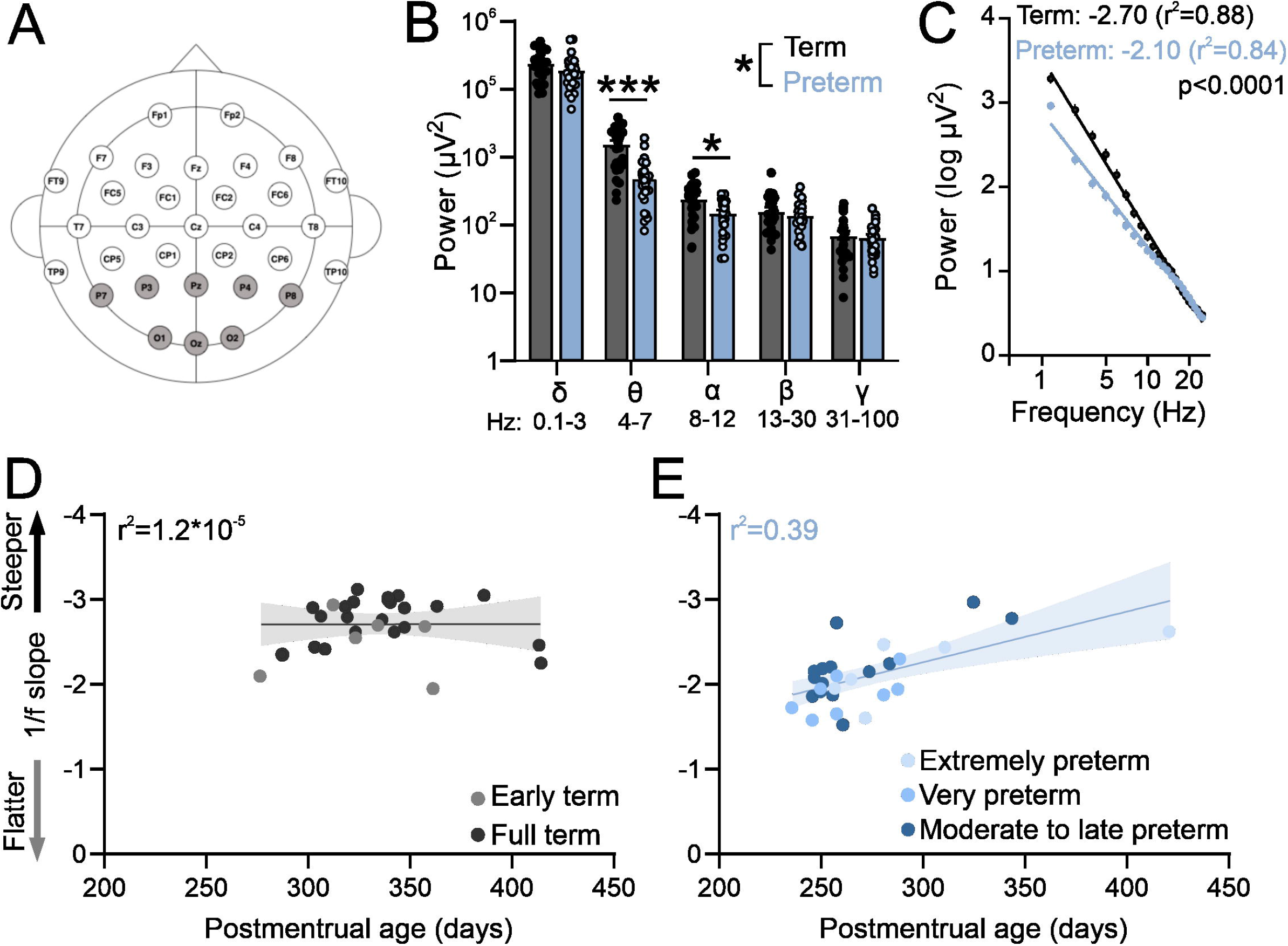
Preterm humans display flatter slope of aperiodic EEG component. **A)** EEG cap montage. Highlighted parietal and occipital channels are analyzed. **B)** Preterm birth significantly reduces resting theta and alpha band power. Two-way ANOVA interaction: p=0.039; F (4, 220) = 2.55; N= 29 preterm and 28 term-born infants. Sidak’s multiple comparisons test: theta p<.0001, alpha p=.022. **C)** Preterm infants have a significantly flatter slope of log transformed 2-25 Hz power. Linear regression; slope and r^2^ values are indicated. Data shown as mean or mean±SEM. **D)** 1/f slope is not correlated with postmenstrual age in term infants. **E)** 1/f slope values are significantly correlated with postmenstrual age in preterm infants (r^2^=0.39, p=0.01).

## Results

Neurocognitive and visual deficits of unclear origin are common in preterm children (Chokron et al., 2021; Macintyre-Béon et al., 2013; Spierer et al., 2004; Edmond and Foroozan, 2006; Weinstein et al., 2012; Emberson et al., 2017), and they are often diagnosed in school age, precluding early interventions (Fazzi et al., 2012; Ben Amor et al., 2012; de Kieviet et al., 2012). To determine how premature birth affects the development of neural activity at a circuit level, we used a cross-species approach to determine the properties of resting activity in a cohort of preterm infants and corrected age-matched term infants, and in a group of preterm mice and their term counterparts.

### Preterm infants and mice display accelerated maturation of resting activity in the visual cortex

We first used EEG in infants to measure resting state activity in the occipital and parietal visual areas within the first 4 months of life. While the preterm infants had a typical distribution of high power in low frequencies and low power in high frequencies (Figure 1 B-C), power in theta and alpha bands was significantly reduced in preterm infants (Figure 1B; Two-way ANOVA frequency x birth interactions F(4,220)=2.55, p=.039; Sidak’s multiple comparison test delta p=.43, theta p<.0001, alpha p=.02, beta p=.95, gamma p=.99).

As visual function matures earlier in preterm infants (Jandó et al., 2012), we then asked if the electrophysiological activity of the visual cortex would reflect this accelerated maturation. To test this, we calculated the slope of aperiodic EEG component 1/f (Figure 1C). 1/f is thought to reflect the background activity of the brain (Gyurkovics et al., 2022; Chini et al., 2022), and 1/f slopes become progressively flatter during infancy (Schaworonkow and Voytek, 2021). As previously reported (Schaworonkow and Voytek, 2021), we found that the power spectra of term and preterm infants were largely aperiodic (Figure 1C). However, we found that preterm infants had a significantly flatter 1/f slope when compared to term infants (F(1,1364)=137.00, p<.0001), despite preterm infants being significantly younger in postmenstrual age (*t*(55)=-6.66, *p*< .001) and equivalent in chronological age (*t*(55)=-0.75, *p*=.455, see also Table 1). 1/f slope was not correlated with postmenstrual age in term infants (Figure 1D), unlike in the preterm cohort, where a significant correlation between postmenstrual age and 1/f slope values was detected (r^2^=.39, p=0.01; Figure 1E). We detected an association (One-way ANOVA, F(4,52)=14.86, p<0.0001) between slope values and the degree of prematurity, with extremely (<28 gestational weeks at birth) and very (28-32 gestational weeks at birth) preterm infants showing the flattest 1/f slopes. We found no effects of mode of delivery on 1/f slope values in either term or preterm cohort (Table 1, C-section vs vaginal t-test; term t(26)=0.19, p=.84; preterm t(27)=0.1, p=.92), nor of respiratory issues reported in both cohorts (Table 1; term, t(26)=0.57, p=0.52; preterm t(27)=0.1, p=0.91). Within the preterm cohort, we further found no relationship between the 1/f slope values and retinopathy of prematurity (t(27)=0.3, p=0.75), or between 1/f slope values and administration of antenatal steroids (t(27)=1.5, p=0.28), two most common complications in our cohort (Table 1). These results confirmed accelerated maturation of visual areas in preterm infants (Jandó et al., 2012; De Asis-Cruz et al., 2020).

To confirm that electrophysiological activity of visual areas in preterm mice recapitulates changes seen in preterm infants (Figure 1C), we used *in vivo* electrophysiology to record intracortical local field potentials (LFPs) in layer 2/3 of the primary visual cortex (V1) of awake, young term and preterm mice (Figure 2E) (Ribic et al., 2019). There was a significant interaction between the timing of birth and the energy composition of the power spectra (Figure 2F; Two-way ANOVA, frequency band x birth interaction F(4,52)=2.93, p=.03), and post-hoc tests revealed a significant reduction of power in the delta band (Figure 2F, Sidak’s multiple comparison test delta p=.001, theta p=.99, alpha, beta and gamma p>.99). However, preterm mice also had a significantly flatter 1/f slope (Figure 2G; F(1,1061)=11.51, p=.0007), indicating accelerated maturation of the primary visual cortex and suggesting the relative conservation of the effects of prematurity on neural activity in mice and humans.

**Figure 2.**
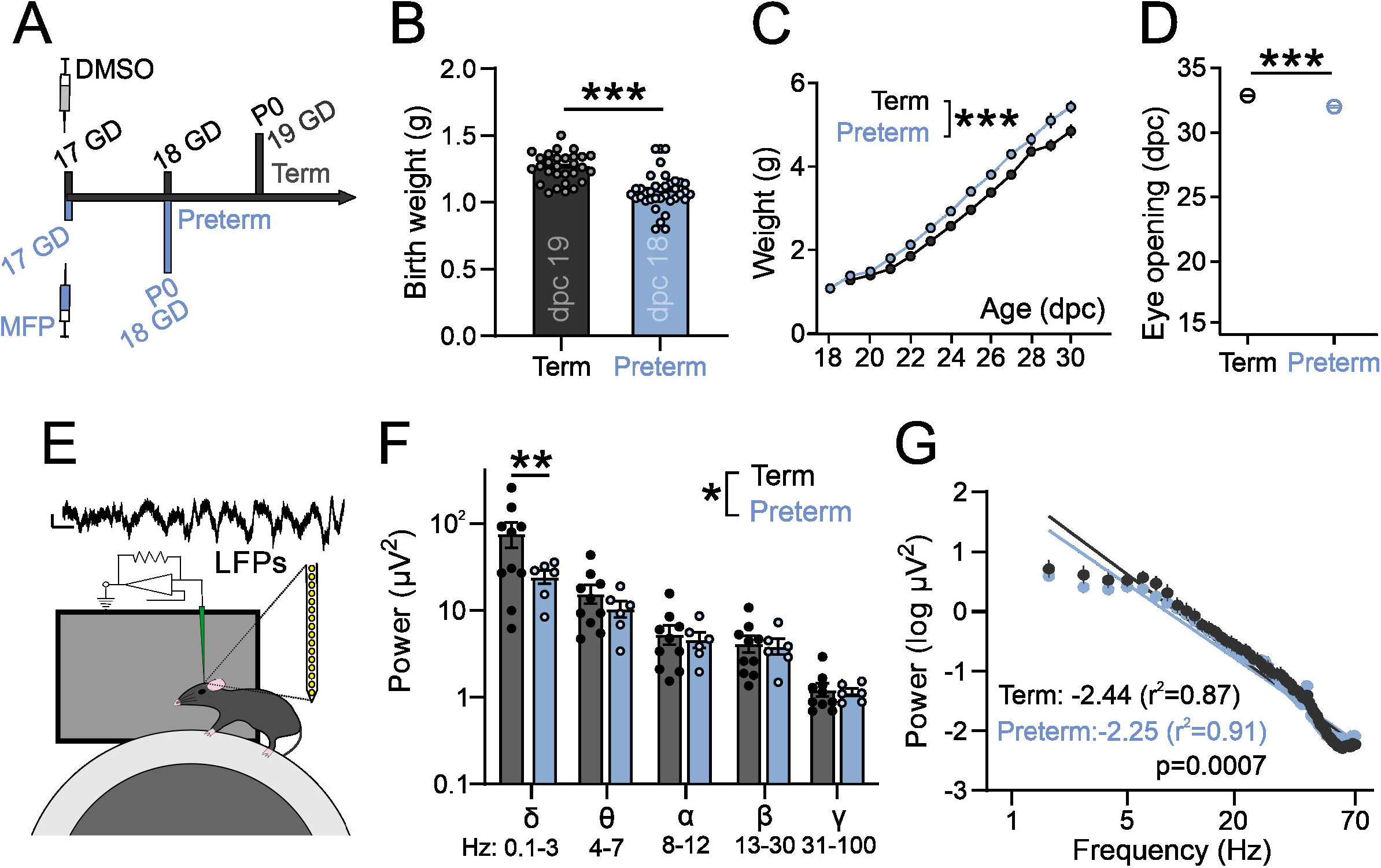
Preterm mice display a flatter slope of aperiodic LFP component. **A)** Preterm mice are generated through the injection of Mifepristone (dissolved in DMSO) to pregnant dams at postconceptional day 17. Dams deliver pups within 24 hrs (1 day early: postnatal day/P 0). Term controls are generated by injecting pregnant dams at postconceptional day 17 with DMSO. **B)** Preterm pups show a significantly lower birth weight (N=29 term and 37 preterm mice, t-test, *t*(64)=5.63, *p*<0.0001) and **C)** an accelerated postnatal growth rate (ordinary two-way ANOVA, *F*(1, 433) = 98.53, *p*<0.0001; N=29 term and 19 preterm mice and **D)** precocious eye opening (t-test, *t*(25)=7.95, *p*<0.0001; N=16 term and 11 preterm mice). **E)** Schematics of in vivo electrophysiology in awake mice. Local field potentials (LFPs) are collected using linear silicone probes (Neuronexus). Scale bar: 10 μV and 0.1 s. **F)** Preterm birth significantly affects the distribution of power across different frequency bands (two-way ANOVA, F(4, 56) = 2.568, p = .048; N=6 preterm and 10 term and mice). G) Preterm mice show a significantly flatter slope of log transformed 2-70 Hz power. Linear regression; slope and r^2^ values are indicated. Data shown as mean or mean±SEM.

### Preterm mice display elevated inhibition in visual cortex

The fourth postnatal week represents a critical transition towards visually-driven activity in mice, reflected in increased suppression of spontaneous neuronal activity by rising levels of inhibitory neurotransmission (Toyoizumi et al., 2013; Shen and Colonnese, 2016). Given the flatter, “older” 1/f slope in preterm mice (Figure 2F-G), we hypothesized that the spontaneous firing rates of visual cortex neurons would be lower in preterm mice reflecting accelerated transition to visually-driven activity. We isolated firing profiles of neurons from all layers of the cortex (Figure 3A) and estimated their firing rates in stationary, awake mice whose eyes were centered on a blank, grey screen (Figure 2E). We indeed found a significantly reduced spontaneous firing rate of visual cortex neurons in preterm mice (Figure 3B; nested t-test t(9)=2.42, p=.030; 239 isolated neurons from 5 term and 6 preterm mice).

**Figure 3.**
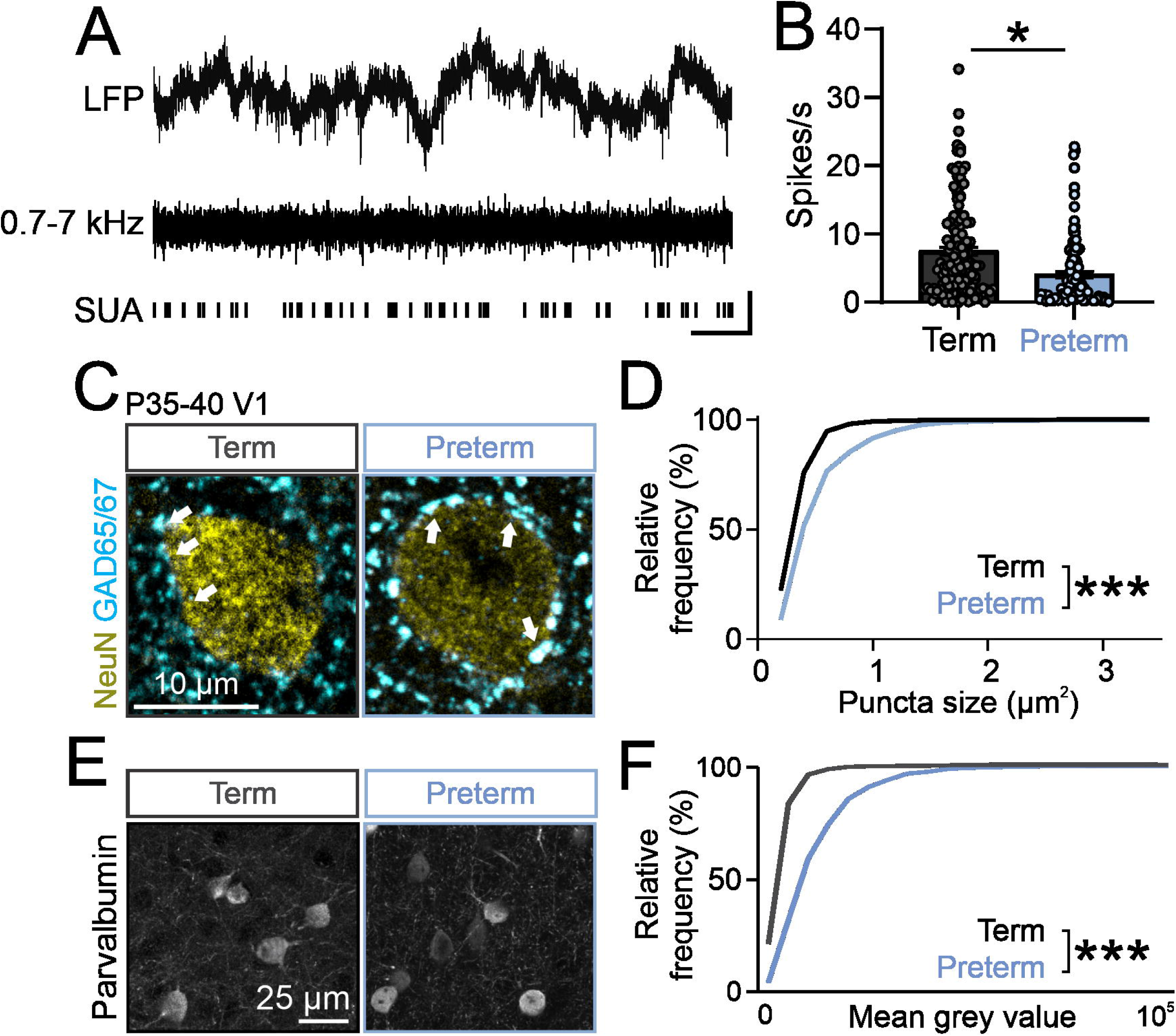
Preterm mice show elevated inhibition in the visual cortex. **A)** Top: raw LFP trace; middle: filtered 0.7-7 kHz LFP; bottom: identified single units. **B)** Preterm mice have a significantly reduced firing rate of neurons in the visual cortex during the presentation of a blank grey screen. 239 units from N=5 term and 6 preterm mice, nested t-test, *t*(9)=2.428, *p*=.038. Data shown as mean or mean±SEM. **C)** Immunohistochemical detection of NeuN (yellow) and GAD65/67 (cyan) in the visual cortex of term and preterm mice. Arrowheads: putative inhibitory synapses. **D)** Cumulative frequency distribution of puncta size in term and preterm mice shows a significant shift to the right in term mice, signifying increased puncta size across all synaptic populations. Kolmogorov-Smirnov test, p<.001. Minimum 100 NeuN cell bodies/mouse from N=5 preterm and 6 term mice. **E)** Representative maximum projections of brain sections stained for Parvalbumin and quantified for Parvalbumin intensity. **F)** Cumulative distribution of Parvalbumin intensity measurements from. The curve in preterm mice is shifter to the right, demonstrating the significantly increased intensity of Parvalbumin staining in preterm mice. N=477 and 636 cell bodies from 3 preterm and 6 term mice, respectively. Kolmogorov-Smirnov test, p<.001.

Suppressed spontaneous firing could be a result of increased inhibition in the developing cortex (Shen and Colonnese, 2016; Toyoizumi et al., 2013; Ribic, 2020). To test this and probe the cellular mechanism of reduced spontaneous firing in preterm mice, we quantified the expression of inhibitory synapse marker glutamic acid decarboxylase 65/67 (GAD65/67) using quantitative immunohistochemistry (Chattopadhyaya et al., 2007; Gogolla et al., 2014; Carcea et al., 2014). GAD65/67 is detected as punctate labeling that surrounds neuronal cell bodies, in line with its localization in the perisomatic inhibitory synapses (Chattopadhyaya et al., 2007). The number and size of perisomatic GAD65/67 puncta are a reliable indicator of inhibitory neurotransmission levels (Carcea et al., 2014). In agreement with suppressed spontaneous activity due to increased inhibition in preterm visual cortex, the size of perisomatic GAD65/67 puncta was significantly increased in preterm mice (Figure 3 E-D; Kolmogorov-Smirnov test p<0.0001, D=0.38), with no changes in their density (Term=15.62±0.4, Preterm=14.98±0.88 puncta/100 μm^2^ of NeuN+ soma; N=6 and 5 mice). Cortical perisomatic inhibition is mediated by fast-spiking, Parvalbumin-expressing interneurons (Pfeffer et al., 2013). As previous studies of preterm birth-related brain injury models reported changes in Parvalbumin interneuron distribution and density (Scheuer et al., 2021; Stolp et al., 2019; Panda et al., 2018), we then asked if this neuronal population is affected by preterm birth. Cortical interneurons represent a mixture of high, middle and low Parvalbumin (PV)-expressing interneurons (Donato et al., 2013), with low PV interneurons being the dominant group in the developing brain and high PV in the mature brain (Donato et al., 2013). In agreement with accelerated maturation of the visual cortex, preterm mice had a significantly shifted distribution of PV intensities, with a higher proportion of middle and high PV-expressing interneurons (Figure 3 E-F; Kolmogorov-Smirnov test D=0.45, p<0.0001), without any changes in the overall density of PV interneurons (Term=22.45±3.53, Preterm=23.79±2.56 PV interneurons per field of view, N=5 term and 3 preterm mice). Altogether, these results demonstrate accelerated maturation of the visual cortex after preterm birth and suggest a central role of inhibition in this process.

## Discussion

Despite extensive research, effects of premature birth on cortical activity in the early postnatal period remain unclear. Through a comparative approach, our study identified accelerated maturation of neural activity in the visual cortex of preterm infants and mice driven by elevated levels of inhibition.

Cortical oscillatory activity shows distinct developmental patterns, with a reduction in the relative power of low frequencies and an increase in high frequencies with increasing age (Gasser et al., 1988; Tierney et al., 2013). Such distribution of powers is likely responsible for the age-related flattening of 1/f EEG slope (Saby and Marshall, 2012; Schaworonkow and Voytek, 2021). An “older” spectral profile in preterm infants and mice is in agreement with the notion that premature exposure to extrauterine environment accelerates brain maturation, at least in primary sensory areas such as the visual cortex (Jandó et al., 2012; De Asis-Cruz et al., 2020). Interestingly, flattened 1/f slope is found in children diagnosed with attention deficit hyperactivity disorder (ADHD), a condition with high prevalence in the preterm population (Arnett et al., 2022; Ostlund et al., 2021; Johnson and Marlow, 2011). Future studies can address if flatter 1/f slope distinguishes preterm-born children with and without ADHD. As primary sensory areas mature earlier than the frontal executive and association areas (Reh et al., 2020; Sydnor et al., 2023), future studies can determine if the accelerated maturation of sensory areas dysregulates the sequence of cortical maturation, impairing the functional maturation of frontal brain areas. This is particularly important given the high prevalence of executive function and visual attention disorders in preterm children (Arpino et al., 2010; Johnson and Marlow, 2011; Burstein et al., 2021).

Fast-spiking, Parvalbumin interneurons are central for the maturation of cortical circuits (Ribic, 2020; Reh et al., 2020). Their functional development is sensitive to experience and in the visual cortex, their maturation can be accelerated or delayed through manipulations of visual input (Ye and Miao, 2013; Guan et al., 2017; Komitova et al., 2013). In agreement, our results suggest that premature onset of visual input can accelerate the maturation of cortical Parvalbumin interneurons, shifting their distribution to mid- and high-PV expressing interneurons in preterm mice. Previous studies of hypoxic mouse models of preterm birth have demonstrated that hypoxia results in reduced density of Parvalbumin interneurons, as well as a decrease in the intensity of Parvalbumin signal (Komitova et al., 2013). The differing findings in our study are likely due to low or absent hypoxia in our mouse model, as well as differences in how preterm brain injury is modelled. Hypoxia models of preterm birth are commonly term-born, with continuous or intermittent exposure to hypoxia during the postnatal development (van der Kooij et al., 2010; Komitova et al., 2013; Salmaso et al., 2014; Lacaille et al., 2019). While hypoxia represents a severe injury, it may not recapitulate the effects of preterm birth alone. Birth itself is an environmental shock that can significantly affect neuronal and synaptic development and accelerate developmental trajectories of different brain areas (Castillo-Ruiz et al., 2020; Toda et al., 2013). As the effects of preterm birth alone on sensory development are unclear, our results highlight a need for multiple animal models to capture the variability in the degree of preterm birth-related brain injury.

Another potential cause of divergence between our findings in preterm mice and previously published findings on interneuronal populations (Lacaille et al., 2019, 2021) in preterm infant cortex is the degree of prematurity. The most vulnerable population of preterm infants are born extremely early, prior to week 28 of gestation, and very early (28-32 weeks of gestation). These are also the infants that show deficits in cortical interneurons (Stolp et al., 2019; Tibrewal et al., 2018; Lacaille et al., 2019, 2021). Yet, the majority (>70%) of preterm infants are born moderately to late preterm (32-37 weeks of gestation), with variable degrees of health complications, including hypoxic brain injury (Karnati et al., 2020; Shapiro-Mendoza and Lackritz, 2012). Infants in our study reflect this, with 51.7% born moderately to late preterm (Table 1). Considering the relatively high viability of preterm mice, as well as our finding of flattest slopes in extremely and very preterm infants, our results suggest that mice born a day early are a model of middle to late preterm birth. However, as both early and late preterm infants have poor neurocognitive outcomes (Karnati et al., 2020; Ben Amor et al., 2012; Crump et al., 2021; Arpino et al., 2010), our study further confirms that birth timing alone is sufficient to dysregulate the maturational trajectory of the brain.

Term-born mouse pups in the first postnatal week are commonly compared to preterm newborns, based on cortical development milestones (Colonnese et al., 2010; Semple et al., 2013). While direct comparison between developmental stages of mice and humans is difficult due to different rates of maturation, our results from the mouse sample confirm previous findings of accelerated brain maturation after premature birth (Castillo-Ruiz et al., 2020; Toda et al., 2013), and provide evidence for it in preterm infants (De Asis-Cruz et al., 2020). Our results further highlight the utility of EEG measures in the clinical setting and set the stage for future longitudinal studies that will explore the relationship between 1/f slope and neurodevelopmental outcomes in the preterm population. Our study adds to the growing body of evidence that birth itself is a critical transition during brain development. Future studies will explore how the timing of this transition affects sensory and cognitive processing, given that preterm infants are at higher risk for developing neurodevelopmental and neuropsychiatric disorders (Johnson and Marlow, 2011).

## Author Contributions

AR and MHP conceived and designed the study. AR acquired and analysed mice data, analysed infant data. and MHP, MN, WC, AB, and AD acquired infant data. MHP analysed infant data. EM analysed infant and mice data. IW, TDH and CS collected and analysed the immunohistochemistry data. AR wrote the manuscript with input from all authors.

## Acknowledgments

We thank the Cang and Liu labs at the Departments of Biology and Psychology for generous access to their confocal microscope, Drs. Zanelli and Fairchild at the University of Virginia Children’s Hospital for identifying candidate participants in the NICU, and the participating families for taking part in our research.

## Data Availability Statement

The datasets generated and analyzed in this study will be provided upon reasonable request.

## Bibliography

Altigani, M., Murphy, J. F., Newcombe, R. G., and Gray, O. P. (1989). Catch up growth in preterm infants. Acta Paediatr. Scand. Suppl. 357, 3–19. doi:10.1111/j.1651-2227.1989.tb11270.x.

Ben Amor, L., Chantal, S., and Bairam, A. (2012). Relationship between late preterm birth and expression of attention-deficit hyperactivity disorder in school-aged children: clinical, neuropsychological, and neurobiochemical outcomes. RRN, 77. doi:10.2147/RRN.S34674.

Antonini, A., Fagiolini, M., and Stryker, M. P. (1999). Anatomical correlates of functional plasticity in mouse visual cortex. J. Neurosci. 19, 4388–4406.

Arnett, A. B., Peisch, V., and Levin, A. R. (2022). The role of aperiodic spectral slope in event-related potentials and cognition among children with and without attention deficit hyperactivity disorder. J. Neurophysiol. 128, 1546–1554. doi:10.1152/jn.00295.2022.

Arpino, C., Compagnone, E., Montanaro, M. L., Cacciatore, D., De Luca, A., Cerulli, A., Di Girolamo, S., and Curatolo, P. (2010). Preterm birth and neurodevelopmental outcome: a review. Childs Nerv. Syst. 26, 1139–1149. doi:10.1007/s00381-010-1125-y.

De Asis-Cruz, J., Kapse, K., Basu, S. K., Said, M., Scheinost, D., Murnick, J., Chang, T., du Plessis, A., and Limperopoulos, C. (2020). Functional brain connectivity in ex utero premature infants compared to in utero fetuses. Neuroimage 219, 117043. doi:10.1016/j.neuroimage.2020.117043.

Ball, G., Srinivasan, L., Aljabar, P., Counsell, S. J., Durighel, G., Hajnal, J. V., Rutherford, M. A., and Edwards, A. D. (2013). Development of cortical microstructure in the preterm human brain. Proc. Natl. Acad. Sci. USA 110, 9541–9546. doi:10.1073/pnas.1301652110.

Barfield, W. D. (2018). Public health implications of very preterm birth. Clin Perinatol 45, 565–577. doi:10.1016/j.clp.2018.05.007.

Blencowe, H., Cousens, S., Oestergaard, M. Z., Chou, D., Moller, A.-B., Narwal, R., Adler, A., Vera Garcia, C., Rohde, S., Say, L., et al. (2012). National, regional, and worldwide estimates of preterm birth rates in the year 2010 with time trends since 1990 for selected countries: a systematic analysis and implications. Lancet 379, 2162–2172. doi:10.1016/S0140-6736(12)60820-4.

Blencowe, H., Lee, A. C. C., Cousens, S., Bahalim, A., Narwal, R., Zhong, N., Chou, D., Say, L., Modi, N., Katz, J., et al. (2013). Preterm birth-associated neurodevelopmental impairment estimates at regional and global levels for 2010. Pediatr. Res. 74 Suppl 1, 17–34. doi:10.1038/pr.2013.204.

Bouyssi-Kobar, M., Brossard-Racine, M., Jacobs, M., Murnick, J., Chang, T., and Limperopoulos, C. (2018). Regional microstructural organization of the cerebral cortex is affected by preterm birth. Neuroimage Clin. 18, 871–880. doi:10.1016/j.nicl.2018.03.020.

Bouyssi-Kobar, M., du Plessis, A. J., McCarter, R., Brossard-Racine, M., Murnick, J., Tinkleman, L., Robertson, R. L., and Limperopoulos, C. (2016). Third trimester brain growth in preterm infants compared with in utero healthy fetuses. Pediatrics 138. doi:10.1542/peds.2016-1640.

Burnsed, J., Skwarzyńska, D., Wagley, P. K., Isbell, L., and Kapur, J. (2019). Neuronal Circuit Activity during Neonatal Hypoxic-Ischemic Seizures in Mice. Ann. Neurol. 86, 927–938. doi:10.1002/ana.25601.

Burstein, O., Zevin, Z., and Geva, R. (2021). Preterm Birth and the Development of Visual Attention During the First 2 Years of Life: A Systematic Review and Meta-analysis. JAMA Netw. Open 4, e213687. doi:10.1001/jamanetworkopen.2021.3687.

Cainelli, E., Vedovelli, L., Wigley, I. L. C. M., Bisiacchi, P. S., and Suppiej, A. (2021). Neonatal spectral EEG is prognostic of cognitive abilities at school age in premature infants without overt brain damage. Eur. J. Pediatr. 180, 909–918. doi:10.1007/s00431-020-03818-x.

Carcea, I., Patil, S. B., Robison, A. J., Mesias, R., Huntsman, M. M., Froemke, R. C., Buxbaum, J. D., Huntley, G. W., and Benson, D. L. (2014). Maturation of cortical circuits requires Semaphorin 7A. Proc. Natl. Acad. Sci. USA 111, 13978–13983. doi:10.1073/pnas.1408680111.

Castillo-Ruiz, A., Hite, T. A., Yakout, D. W., Rosen, T. J., and Forger, N. G. (2020). Does birth trigger cell death in the developing brain? eNeuro 7. doi:10.1523/ENEURO.0517-19.2020.

Chattopadhyaya, B., Di Cristo, G., Wu, C. Z., Knott, G., Kuhlman, S., Fu, Y., Palmiter, R. D., and Huang, Z. J. (2007). GAD67-mediated GABA synthesis and signaling regulate inhibitory synaptic innervation in the visual cortex. Neuron 54, 889–903. doi:10.1016/j.neuron.2007.05.015.

Chiesa, M., Guimond, D., Tyzio, R., Pons-Bennaceur, A., Lozovaya, N., Burnashev, N., Ferrari, D. C., and Ben-Ari, Y. (2019). Term or Preterm Cesarean Section Delivery Does Not Lead to Long-term Detrimental Consequences in Mice. Cereb. Cortex 29, 2424–2436. doi:10.1093/cercor/bhy112.

Chini, M., Pfeffer, T., and Hanganu-Opatz, I. (2022). An increase of inhibition drives the developmental decorrelation of neural activity. Elife 11. doi:10.7554/eLife.78811.

Chokron, S., Kovarski, K., and Dutton, G. N. (2021). Cortical visual impairments and learning disabilities. Front. Hum. Neurosci. 15, 713316. doi:10.3389/fnhum.2021.713316.

Colonnese, M. T., Kaminska, A., Minlebaev, M., Milh, M., Bloem, B., Lescure, S., Moriette, G., Chiron, C., Ben-Ari, Y., and Khazipov, R. (2010). A conserved switch in sensory processing prepares developing neocortex for vision. Neuron 67, 480–498. doi:10.1016/j.neuron.2010.07.015.

Crump, C., Sundquist, J., and Sundquist, K. (2021). Preterm or early term birth and risk of autism. Pediatrics 148. doi:10.1542/peds.2020-032300.

de Kieviet, J. F., van Elburg, R. M., Lafeber, H. N., and Oosterlaan, J. (2012). Attention problems of very preterm children compared with age-matched term controls at school-age. J. Pediatr. 161, 824–829. doi:10.1016/j.jpeds.2012.05.010.

Donato, F., Rompani, S. B., and Caroni, P. (2013). Parvalbumin-expressing basket-cell network plasticity induced by experience regulates adult learning. Nature 504, 272–276. doi:10.1038/nature12866.

Dudley, D. J., Branch, D. W., Edwin, S. S., and Mitchell, M. D. (1996). Induction of preterm birth in mice by RU486. Biol. Reprod. 55, 992–995. doi:10.1095/biolreprod55.5.992.

Edmond, J. C., and Foroozan, R. (2006). Cortical visual impairment in children. Curr Opin Ophthalmol 17, 509–512. doi:10.1097/ICU.0b013e3280107bc5.

El-Hayek, Y. H., Wu, C., and Zhang, L. (2011). Early suppression of intracranial EEG signals predicts ischemic outcome in adult mice following hypoxia-ischemia. Exp. Neurol. 231, 295–303. doi:10.1016/j.expneurol.2011.07.003.

Elovitz, M. A., and Mrinalini, C. (2004). Animal models of preterm birth. Trends Endocrinol. Metab. 15, 479–487. doi:10.1016/j.tem.2004.10.009.

Emberson, L. L., Boldin, A. M., Riccio, J. E., Guillet, R., and Aslin, R. N. (2017). Deficits in Top-Down Sensory Prediction in Infants At Risk due to Premature Birth. Curr. Biol. 27, 431–436. doi:10.1016/j.cub.2016.12.028.

Fawcett, S. L., Wang, Y.-Z., and Birch, E. E. (2005). The critical period for susceptibility of human stereopsis. Invest. Ophthalmol. Vis. Sci. 46, 521–525. doi:10.1167/iovs.04-0175.

Fazzi, E., Galli, J., and Micheletti, S. (2012). Visual impairment: A common sequela of preterm birth. Neoreviews 13, e542–e550. doi:10.1542/neo.13-9-e542.

Freeman, R. D., and Ohzawa, I. (1992). Development of binocular vision in the kitten’s striate cortex. J. Neurosci. 12, 4721–4736. doi:10.1523/JNEUROSCI.12-12-04721.1992.

Gasser, T., Verleger, R., Bächer, P., and Sroka, L. (1988). Development of the EEG of school-age children and adolescents. I. Analysis of band power. Electroencephalogr. Clin. Neurophysiol. 69, 91–99. doi:10.1016/0013-4694(88)90204-0.

Gogolla, N., Takesian, A. E., Feng, G., Fagiolini, M., and Hensch, T. K. (2014). Sensory integration in mouse insular cortex reflects GABA circuit maturation. Neuron 83, 894–905. doi:10.1016/j.neuron.2014.06.033.

Guan, W., Cao, J.-W., Liu, L.-Y., Zhao, Z.-H., Fu, Y., and Yu, Y.-C. (2017). Eye opening differentially modulates inhibitory synaptic transmission in the developing visual cortex. Elife 6. doi:10.7554/eLife.32337.

Gyurkovics, M., Clements, G. M., Low, K. A., Fabiani, M., and Gratton, G. (2022). Stimulus-induced changes in 1/f-like background activity in EEG. J. Neurosci. 42, 7144–7151. doi:10.1523/JNEUROSCI.0414-22.2022.

Haak, K. V., Morland, A. B., and Engel, S. A. (2015). Plasticity, and its limits, in adult human primary visual cortex. Multisens. Res. 28, 297–307. doi:10.1163/22134808-00002496.

Holmes, G. L., and Lombroso, C. T. (1993). Prognostic value of background patterns in the neonatal EEG. J Clin Neurophysiol 10, 323–352.

Del Hoyo Soriano, L., Rosser, T., Hamilton, D., Wood, T., Abbeduto L., and Sherman, S. (2020). Gestational age is related to symptoms of attention-deficit/hyperactivity disorder in late-preterm to full-term children and adolescents with down syndrome. Sci. Rep. 10, 20345. doi:10.1038/s41598-020-77392-5.

Hunt, B. A. E., Scratch, S. E., Mossad, S. I., Emami, Z., Taylor, M. J., and Dunkley, B. T. (2019). Disrupted visual cortex neurophysiology following very preterm birth. Biol. Psychiatry Cogn. Neurosci. Neuroimaging. doi:10.1016/j.bpsc.2019.08.012.

Huttenlocher, P. R., de Courten, C., Garey, L. J., and Van der Loos, H. (1982). Synaptogenesis in human visual cortex--evidence for synapse elimination during normal development. Neurosci. Lett. 33, 247–252. doi:10.1016/0304-3940(82)90379-2.

Jaekel, J., Wolke, D., and Bartmann, P. (2013). Poor attention rather than hyperactivity/impulsivity predicts academic achievement in very preterm and full-term adolescents. Psychol. Med. 43, 183–196. doi:10.1017/S0033291712001031.

Jakab, A., Natalucci, G., Koller, B., Tuura, R., Rüegger, C., and Hagmann, C. (2020). Mental development is associated with cortical connectivity of the ventral and nonspecific thalamus of preterm newborns. Brain Behav., e01786. doi:10.1002/brb3.1786.

Jandó, G., Mikó-Baráth, E., Markó, K., Hollódy, K., Török, B., and Kovacs, I. (2012). Early-onset binocularity in preterm infants reveals experience-dependent visual development in humans. Proc. Natl. Acad. Sci. USA 109, 11049–11052. doi:10.1073/pnas.1203096109.

Johnson, K. J., Moy, B., Rensing, N., Robinson, A., Ly, M., Chengalvala, R., Wong, M., and Galindo, R. (2022). Functional neuropathology of neonatal hypoxia-ischemia by single-mouse longitudinal electroencephalography. Epilepsia 63, 3037–3050. doi:10.1111/epi.17403.

Johnson, S., and Marlow, N. (2011). Preterm birth and childhood psychiatric disorders. Pediatr. Res. 69, 11R–8R. doi:10.1203/PDR.0b013e318212faa0.

Karnati, S., Kollikonda, S., and Abu-Shaweesh, J. (2020). Late preterm infants - Changing trends and continuing challenges. Int. J. Pediatr. Adolesc. Med. 7, 36–44. doi:10.1016/j.ijpam.2020.02.006.

Kiorpes, L. (2016). The puzzle of visual development: behavior and neural limits. J. Neurosci. 36, 11384–11393. doi:10.1523/JNEUROSCI.2937-16.2016.

Komitova, M., Xenos, D., Salmaso, N., Tran, K. M., Brand, T., Schwartz, M. L., Ment, L., and Vaccarino, F. M. (2013). Hypoxia-induced developmental delays of inhibitory interneurons are reversed by environmental enrichment in the postnatal mouse forebrain. J. Neurosci. 33, 13375–13387. doi:10.1523/JNEUROSCI.5286-12.2013.

Kozhemiako, N., Nunes, A., Vakorin, V. A., Chau, C. M. Y., Moiseev, A., Ribary, U., Grunau, R. E., and Doesburg, S. M. (2019). Atypical resting state neuromagnetic connectivity and spectral power in very preterm children. J. Child Psychol. Psychiatry 60, 975–987. doi:10.1111/jcpp.13026.

Lacaille, H., Vacher, C.-M., Bakalar, D., O’Reilly, J. J., Salzbank, J., and Penn, A. A. (2019). Impaired interneuron development in a novel model of neonatal brain injury. eNeuro 6. doi:10.1523/ENEURO.0300-18.2019.

Lacaille, H., Vacher, C.-M., and Penn, A. A. (2021). Preterm birth alters the maturation of the gabaergic system in the human prefrontal cortex. Front. Mol. Neurosci. 14, 827370. doi:10.3389/fnmol.2021.827370.

Leung, M. P., Thompson, B., Black, J., Dai, S., and Alsweiler, J. M. (2018). The effects of preterm birth on visual development. Clin Exp Optom 101, 4–12. doi:10.1111/cxo.12578.

Limperopoulos, C., Bassan, H., Gauvreau, K., Robertson, R. L., Sullivan, N. R., Benson, C. B., Avery, L., Stewart, J., Soul, J. S., Ringer, S. A., et al. (2007). Does cerebellar injury in premature infants contribute to the high prevalence of long-term cognitive, learning, and behavioral disability in survivors? Pediatrics 120, 584–593. doi:10.1542/peds.2007-1041.

Lloyd, R. O., O’Toole, J. M., Livingstone, V., Filan, P. M., and Boylan, G. B. (2021). Can EEG accurately predict 2-year neurodevelopmental outcome for preterm infants? Arch. Dis. Child. Fetal Neonatal Ed. 106, 535–541. doi:10.1136/archdischild-2020-319825.

Lunghi, C., Burr, D. C., and Morrone, C. (2011). Brief periods of monocular deprivation disrupt ocular balance in human adult visual cortex. Curr. Biol. 21, R538–9. doi:10.1016/j.cub.2011.06.004.

Macintyre-Béon, C., Young, D., Dutton, G. N., Mitchell, K., Simpson, J., Loffler, G., Bowman, R., and Hamilton, R. (2013). Cerebral visual dysfunction in prematurely born children attending mainstream school. Doc. Ophthalmol. 127, 89–102. doi:10.1007/s10633-013-9405-y.

Malik, S., Vinukonda, G., Vose, L. R., Diamond, D., Bhimavarapu, B. B. R., Hu, F., Zia, M. T., Hevner, R., Zecevic, N., and Ballabh, P. (2013). Neurogenesis continues in the third trimester of pregnancy and is suppressed by premature birth. J. Neurosci. 33, 411–423. doi:10.1523/JNEUROSCI.4445-12.2013.

Mann, J. R., McDermott, S., Griffith, M. I., Hardin, J., and Gregg, A. (2011). Uncovering the complex relationship between pre-eclampsia, preterm birth and cerebral palsy. Paediatr Perinat Epidemiol 25, 100–110. doi:10.1111/j.1365-3016.2010.01157.x.

McCarthy, R., Martin-Fairey, C., Sojka, D. K., Herzog, E. D., Jungheim, E. S., Stout, M. J., Fay, J. C., Mahendroo, M., Reese, J., Herington, J. L., et al. (2018). Mouse models of preterm birth: suggested assessment and reporting guidelines. Biol. Reprod. 99, 922–937. doi:10.1093/biolre/ioy109.

McCoy, B., and Hahn, C. D. (2013). Continuous EEG monitoring in the neonatal intensive care unit. J Clin Neurophysiol 30, 106–114. doi:10.1097/WNP.0b013e3182872919.

McGowan, E. C., and Sheinkopf, S. J. (2021). Autism and preterm birth: clarifying risk and exploring mechanisms. Pediatrics 148. doi:10.1542/peds.2021-051978.

Mitchell, D. E., and Maurer, D. (2022). Critical periods in vision revisited. Annu. Rev. Vis. Sci. 8, 291–321. doi:10.1146/annurev-vision-090721-110411.

Nishiyori, R., Xiao, R., Vanderbilt, D., and Smith, B. A. (2021). Electroencephalography measures of relative power and coherence as reaching skill emerges in infants born preterm. Sci. Rep. 11, 3609. doi:10.1038/s41598-021-82329-7.

Norcia, A. M., Tyler, C. W., Piecuch, R., Clyman, R., and Grobstein, J. (1987). Visual acuity development in normal and abnormal preterm human infants. J Pediatr Ophthalmol Strabismus 24, 70–74.

Nordvik, T., Schumacher, E. M., Larsson, P. G., Pripp, A. H., Løhaugen, G. C., and Stiris, T. (2022). Early spectral EEG in preterm infants correlates with neurocognitive outcomes in late childhood. Pediatr. Res. 92, 1132–1139. doi:10.1038/s41390-021-01915-7.

O’Toole, J. M., and Boylan, G. B. (2019). Quantitative preterm EEG analysis: the need for caution in using modern data science techniques. Front. Pediatr. 7, 174. doi:10.3389/fped.2019.00174.

Ortinau, C., and Neil, J. (2015). The neuroanatomy of prematurity: normal brain development and the impact of preterm birth. Clin Anat 28, 168–183. doi:10.1002/ca.22430.

Ostlund, B. D., Alperin, B. R., Drew, T., and Karalunas, S. L. (2021). Behavioral and cognitive correlates of the aperiodic (1/f-like) exponent of the EEG power spectrum in adolescents with and without ADHD. Dev Cogn Neurosci 48, 100931. doi:10.1016/j.dcn.2021.100931.

Panda, S., Dohare, P., Jain, S., Parikh, N., Singla, P., Mehdizadeh, R., Klebe, D. W., Kleinman, G. M., Cheng, B., and Ballabh, P. (2018). Estrogen Treatment Reverses Prematurity-Induced Disruption in Cortical Interneuron Population. J. Neurosci. 38, 7378–7391. doi:10.1523/JNEUROSCI.0478-18.2018.

Pandit, A. S., Robinson, E., Aljabar, P., Ball, G., Gousias, I. S., Wang, Z., Hajnal, J. V., Rueckert, D., Counsell, S. J., Montana, G., et al. (2014). Whole-brain mapping of structural connectivity in infants reveals altered connection strength associated with growth and preterm birth. Cereb. Cortex 24, 2324–2333. doi:10.1093/cercor/bht086.

Pfeffer, C. K., Xue, M., He, M., Huang, Z. J., and Scanziani, M. (2013). Inhibition of inhibition in visual cortex: the logic of connections between molecularly distinct interneurons. Nat. Neurosci. 16, 1068–1076. doi:10.1038/nn.3446.

Pinto, J. G. A., Hornby, K. R., Jones, D. G., and Murphy, K. M. (2010). Developmental changes in GABAergic mechanisms in human visual cortex across the lifespan. Front. Cell Neurosci. 4, 16. doi:10.3389/fncel.2010.00016.

Reh, R. K., Dias, B. G., Nelson, C. A., Kaufer, D., Werker, J. F., Kolb, B., Levine, J. D., and Hensch, T. K. (2020). Critical period regulation across multiple timescales. Proc. Natl. Acad. Sci. USA 117, 23242–23251. doi:10.1073/pnas.1820836117.

Ribic, A. (2020). Stability in the Face of Change: Lifelong Experience-Dependent Plasticity in the Sensory Cortex. Front. Cell Neurosci. 14, 76. doi:10.3389/fncel.2020.00076.

Ribic, A., Crair, M. C., and Biederer, T. (2019). Synapse-Selective Control of Cortical Maturation and Plasticity by Parvalbumin-Autonomous Action of SynCAM 1. Cell Rep. 26, 381–393.e6. doi:10.1016/j.celrep.2018.12.069.

Rivera, M. J., Teruel, M. A., Maté, A., and Trujillo, J. (2021). Diagnosis and prognosis of mental disorders by means of EEG and deep learning: a systematic mapping study. Artif. Intell. Rev. doi:10.1007/s10462-021-09986-y.

Saby, J. N., and Marshall, P. J. (2012). The utility of EEG band power analysis in the study of infancy and early childhood. Dev. Neuropsychol. 37, 253–273. doi:10.1080/87565641.2011.614663.

Salmaso, N., Jablonska, B., Scafidi, J., Vaccarino, F. M., and Gallo, V. (2014). Neurobiology of premature brain injury. Nat. Neurosci. 17, 341–346. doi:10.1038/nn.3604.

Schaworonkow, N., and Voytek, B. (2021). Longitudinal changes in aperiodic and periodic activity in electrophysiological recordings in the first seven months of life. Dev Cogn Neurosci 47, 100895. doi:10.1016/j.dcn.2020.100895.

Scheuer, T., dem Brinke, E. A., Grosser, S., Wolf, S. A., Mattei, D., Sharkovska, Y., Barthel, P. C., Endesfelder, S., Friedrich, V., Bührer, C., et al. (2021). Reduction of cortical parvalbumin-expressing GABAergic interneurons in a rodent hyperoxia model of preterm birth brain injury with deficits in social behavior and cognition. Development 148. doi:10.1242/dev.198390.

Schwindt, E., Giordano, V., Rona, Z., Czaba-Hnizdo, C., Olischar, M., Waldhoer, T., Werther, T., Fuiko, R., Berger, A., and Klebermass-Schrehof, K. (2018). The impact of extrauterine life on visual maturation in extremely preterm born infants. Pediatr. Res. 84, 403–410. doi:10.1038/s41390-018-0084-y.

Semple, B. D., Blomgren, K., Gimlin, K., Ferriero, D. M., and Noble-Haeusslein, L. J. (2013). Brain development in rodents and humans: Identifying benchmarks of maturation and vulnerability to injury across species. Prog. Neurobiol. 106-107, 1–16. doi:10.1016/j.pneurobio.2013.04.001.

Shapiro-Mendoza, C. K., and Lackritz, E. M. (2012). Epidemiology of late and moderate preterm birth. Semin Fetal Neonatal Med 17, 120–125. doi:10.1016/j.siny.2012.01.007.

Shen, J., and Colonnese, M. T. (2016). Development of activity in the mouse visual cortex. J. Neurosci. 36, 12259–12275. doi:10.1523/JNEUROSCI.1903-16.2016.

Shuffrey, L. C., Pini, N., Potter, M., Springer, P., Lucchini, M., Rayport, Y., Sania, A., Firestein, M., Brink, L., Isler, J. R., et al. (2022). Aperiodic electrophysiological activity in preterm infants is linked to subsequent autism risk. Dev. Psychobiol. 64, e22271. doi:10.1002/dev.22271.

Spierer, A., Royzman, Z., and Kuint, J. (2004). Visual acuity in premature infants. Ophthalmologica 218, 397–401. doi:10.1159/000080943.

Stolp, H. B., Fleiss, B., Arai, Y., Supramaniam, V., Vontell, R., Birtles, S., Yates, A. G., Baburamani, A. A., Thornton, C., Rutherford, M., et al. (2019). Interneuron development is disrupted in preterm brains with diffuse white matter injury: observations in mouse and human. Front. Physiol. 10, 955. doi:10.3389/fphys.2019.00955.

Strang-Karlsson, S., Andersson, S., Paile-Hyvärinen, M., Darby, D., Hovi, P., Räikkönen, K., Pesonen, A.-K., Heinonen, K., Järvenpää, A.-L., Eriksson, J. G., et al. (2010). Slower reaction times and impaired learning in young adults with birth weight <1500 g. Pediatrics 125, e74–82. doi:10.1542/peds.2009-1297.

Sydnor, V. J., Larsen, B., Seidlitz, J., Adebimpe, A., Alexander-Bloch, A. F., Bassett, D. S., Bertolero, M. A., Cieslak, M., Covitz, S., Fan, Y., et al. (2023). Intrinsic activity development unfolds along a sensorimotor–association cortical axis in youth. Nat. Neurosci. doi:10.1038/s41593-023-01282-y.

Tibrewal, M., Cheng, B., Dohare, P., Hu, F., Mehdizadeh, R., Wang, P., Zheng, D., Ungvari, Z., and Ballabh, P. (2018). Disruption of Interneuron Neurogenesis in Premature Newborns and Reversal with Estrogen Treatment. J. Neurosci. 38, 1100–1113. doi:10.1523/JNEUROSCI.1875-17.2017.

Tierney, A., Strait, D. L., O’Connell, S., and Kraus, N. (2013). Developmental changes in resting gamma power from age three to adulthood. Clin. Neurophysiol. 124, 1040–1042. doi:10.1016/j.clinph.2012.09.023.

Toda, T., Homma, D., Tokuoka, H., Hayakawa, I., Sugimoto, Y., Ichinose, H., and Kawasaki, H. (2013). Birth regulates the initiation of sensory map formation through serotonin signaling. Dev. Cell 27, 32–46. doi:10.1016/j.devcel.2013.09.002.

Tokariev, A., Stjerna, S., Lano, A., Metsäranta, M., Palva, J. M., and Vanhatalo, S. (2019). Preterm birth changes networks of newborn cortical activity. Cereb. Cortex 29, 814–826. doi:10.1093/cercor/bhy012.

Toyoizumi, T., Miyamoto, H., Yazaki-Sugiyama, Y., Atapour, N., Hensch, T. K., and Miller, K. D. (2013). A theory of the transition to critical period plasticity: inhibition selectively suppresses spontaneous activity. Neuron 80, 51–63. doi:10.1016/j.neuron.2013.07.022.

Treyvaud, K., Ure, A., Doyle, L. W., Lee, K. J., Rogers, C. E., Kidokoro, H., Inder, T. E., and Anderson, P. J. (2013). Psychiatric outcomes at age seven for very preterm children: rates and predictors. J. Child Psychol. Psychiatry 54, 772–779. doi:10.1111/jcpp.12040.

van der Kooij, M. A., Ohl, F., Arndt, S. S., Kavelaars, A., van Bel, F., and Heijnen, C. J. (2010). Mild neonatal hypoxia-ischemia induces long-term motor- and cognitive impairments in mice. Brain Behav. Immun. 24, 850–856. doi:10.1016/j.bbi.2009.09.003.

Vanhatalo, S., Tallgren, P., Andersson, S., Sainio, K., Voipio, J., and Kaila, K. (2002). DC-EEG discloses prominent, very slow activity patterns during sleep in preterm infants. Clin. Neurophysiol. 113, 1822–1825. doi:10.1016/S1388-2457(02)00292-4.

Wang, B.-S., Sarnaik, R., and Cang, J. (2010). Critical period plasticity matches binocular orientation preference in the visual cortex. Neuron 65, 246–256. doi:10.1016/j.neuron.2010.01.002.

Watanabe, K., Hayakawa, F., and Okumura, A. (1999). Neonatal EEG: a powerful tool in the assessment of brain damage in preterm infants. Brain and Development 21, 361–372. doi:10.1016/S0387-7604(99)00034-0.

Wehrle, F. M., Michels, L., Guggenberger, R., Huber, R., Latal, B., O’Gorman, R. L., and Hagmann, C. F. (2018). Altered resting-state functional connectivity in children and adolescents born very preterm short title. Neuroimage Clin. 20, 1148–1156. doi:10.1016/j.nicl.2018.10.002.

Weinstein, J. M., Gilmore, R. O., Shaikh, S. M., Kunselman, A. R., Trescher, W. V., Tashima, L. M., Boltz, M. E., McAuliffe, M. B., Cheung, A., and Fesi, J. D. (2012). Defective motion processing in children with cerebral visual impairment due to periventricular white matter damage. Dev. Med. Child Neurol. 54, e1–8. doi:10.1111/j.1469-8749.2010.03874.x.

Ye, Q., and Miao, Q.-L. (2013). Experience-dependent development of perineuronal nets and chondroitin sulfate proteoglycan receptors in mouse visual cortex. Matrix Biol 32, 352–363. doi:10.1016/j.matbio.2013.04.001.

Zanelli, S., Goodkin, H. P., Kowalski, S., and Kapur, J. (2014). Impact of transient acute hypoxia on the developing mouse EEG. Neurobiol. Dis. 68, 37–46. doi:10.1016/j.nbd.2014.03.005.

