## Supplemental Materials for "Preterm birth accelerates the maturation of spontaneous and resting activity in the visual cortex"

**Supplementary Materials**

**Method**

*Infant EEG acquisition and preprocessing*.

Resting-state EEG was recorded from 32 Ag/AgCl active actiCAP slim electrodes (Brain Products GmbH, Germany) affixed to an elastic cap according to the 10–20 electrode placement system (Figure 1A) while the infant rested in a caregiver’s arms for up to 7 min. EEG was amplified with a BrainAmp DC Amplifier and recorded using BrainVision Recorder software with a sampling rate of 5000 Hz, online referenced to FCz, and online band-pass filtered between 0.01 and 1000 Hz. Data were preprocessed with an automated preprocessing pipeline specifically validated and shown to generate reliable estimates for the computation of aperiodic signal in pediatric EEG data (APPLESED, Puglia et al., 2022) using EEGLab v2021.1 software with quality assurance via manual inspection: data were down-sampled to 500 Hz, band-pass filtered from 0.1-100 Hz, and segmented into 10s epochs (Delorme and Makeig, 2004). Epochs with a voltage exceeding ± 500 µV were rejected. The data was then decomposed via independent components analysis and artifactual components were removed using the adjusted-ADJUST algorithm optimized for use in infant data (Debnath et al., 2020; Mognon et al., 2011). The number of components removed did not significantly differ across preterm (M=5.75) and term (M=5.89) groups (t(55)=-0.15, p=.883). Epochs with amplitude standard deviations exceeding 80 μV within a 200-ms sliding window with a 100-ms window step were discarded and problematic channels were interpolated (Nolan et al., 2010). The number of channels interpolated ranged from 0 to 3 and did not significantly differ across the preterm (M=0.93) and term (M=0.71) groups (t(55)=0.96, p=.339). Finally, the 6 epochs with a total global field power (GFP) closest to the median GFP for each participant were selected for spectral analysis. Thirty-eight preterm and 30 term infants had sufficient, artifact-free data after preprocessing. An additional 11 infants (9 preterm) were excluded due to high line noise (60 Hz) contamination and mean signal amplitude < 1 μV. See Table 1 for participant demographic and perinatal characteristics for the final sample.

*In vivo electrophysiology in mice.* Recordings were performed on awake term and preterm (N=9 and 6, respectively) female and male mice, ages 21 to 28 days after birth, using a treadmill as described in (Niell and Stryker, 2010). Four to seven days before the recording session, custom made stainless steel head-plate implants were cemented to the mouse skull to stabilize the head and affix the animal to the recording setup. Animals were anesthetized with isoflurane in oxygen (2% induction, 1.0–1.8% maintenance), warmed with a heating pad at 38°C and given subcutaneous injections of Buprenorphine SR (1mg/kg) and 0.25% Bupivacaine (locally) for analgesia. Eyes were covered with Puralube (Decra, Northwich, UK) to prevent drying. Scalp and fascia from Bregma to behind lambda were removed, and the skull was cleaned, dried and covered with a thin layer of Scotchbond adhesive (3M, Maplewood, MN) as a cement primer. Skin edges were sealed with VetBond (3M). The head plate was attached with dental cement (RelyX Ultimate, 3M). The well of the head plate was filled with silicone elastomer (Reynold Advanced Materials, Brighton, MA) to protect the skull before recordings. Animals were group housed after the implantation and monitored daily for signs of shock or infection. Two to three days before the recording, the animals underwent one to two 20-30 minutes handling sessions and one to two 10-20 minutes sessions in which the animals were habituated to the treadmill (Dombeck et al., 2007). On the day of recording, the animals were anesthetized as above and small craniotomies (~0.5 mm in diameter) with 18G needles were made above V1 (2-3 mm lateral to midline, 0.5-1 mm anterior to lambda) and cerebellum. The brain surface was covered in 2-3% low melting point agarose (Promega, Madison, WI) in sterile saline and then capped with silicone elastomer. Animals were allowed to recover for 2–4 h. For the recording sessions, mice were placed in the head-plate holder above the treadmill and allowed to habituate for 5-10 minutes. The agarose and silicone plug were removed, the reference insulated silver wire electrode (A-M Systems, Carlsborg, WA) was placed in cerebellum and the well was covered with warm sterile saline. A multisite electrode spanning all cortical layers (A1x16-5mm-50-177-A16; Neuronexus Technologies, Ann Arbor, MI) was coated with DiI (Invitrogen) to allow post hoc insertion site verification and then inserted in the brain through the craniotomy. The electrode was lowered until the uppermost recording site had entered the brain and allowed to settle for 20-30 minutes. The well with the electrode was then filled with 3% agarose to stabilize the electrode and the whole region was kept moist with surgical gelfoam soaked in sterile saline (Pfizer, MA). Minimum 2 penetrations were made per animal to ensure proper sampling of the craniotomy. After the recording, mice were euthanized with an overdose of ketamine and xylazine or kept for subsequent experiments after protecting the craniotomy with silicone elastomer.

*Data collection and analysis for mice.* Blank screen was generated with MATLAB (MathWorks, Natick, MA) using the Psychtoolbox extension (Brainard, 1997) and presented on a gamma corrected 27” LCD. The screen was centered 25 cm from the mouse's eye, covering ∼80° of visual space. The signals were sampled at 25 kHz using Spike2 and data acquisition unit (Power 1401-3, CED). Signals were fed into a 16-channel amplifier (Model 3500; A-M Systems), amplified 200x and band-pass filtered 0.7-7000 Hz. Only stationary, non-running stages were analyzed offline using Spike2 software (CED). For single unit analysis, spikes were extracted from band-pass filtered data (all 16 channels) using thresholds (3x standard deviation) and sorted in Spike 2. For spectral analyses, layer 2/3 was selected due to its high correlation with EEG signal (Goswami et al., 2019). 60 s epochs of data during viewing of the blank screen were selected and downsampled to 500 Hz prior to spectral analysis to match the infant data.

*Immunohistochemistry and imaging.* Term and preterm mice (aged 35-40 days, N=3-6/group, as indicated in text and figure legends) were anesthetized with a mixture of ketamine and xylazine and transcranially perfused with warm 0.1 M phosphate buffer, followed by warm 4% paraformaldehyde (Electron Microscopy Sciences, Hatfield, PA). Brains were postfixed 1 hr at room temperature, followed by overnight fixation at 4°C. Brains were sectioned into 40 µm sections using a vibratome and stored in 1x phosphate buffered saline (PBS) and 0.01% sodium azide. For immunohistochemistry, sections were rinsed in PBS, non-specific binding was blocked with 3% normal horse serum (heat inactivated, ThermoFisher, Waltham, MA) and 0.3% Triton-X 100 (Sigma-Aldrich) in PBS (sterile filtered). Antibodies were incubated overnight at 4°C. Parvalbumin (anti-goat) was used at 1/200 (Swant, Belinzona, Switzerland) and detected with donkey anti-goat Alexa 647 (ThermoFisher). NeuN and GAD65/67 were both anti-rabbit (Milipore Sigma) and were used with secondary NanoTag reagents (FluoTag X2 Atto 488 and Alexa 647) according to the manufacturer’s protocol (NanoTag Biotechnologies, GmbH, Goettingen, Germany). After staining, sections were rinsed in distilled water, mounted on glass slides, briefly dried, and coverslipped with Aquamount (Polysciences, Warrington, PA). Images were acquired using Zeiss LSM 800 at 2048x2048 resolution. Single optical sections from the visual cortex using 63x 1.2 NA C-Apochromat were acquired for GAD65/67 quantification, and z-stacks were acquired using 40x1.2 NA Plan-Apochromat for Parvalbumin intensity quantification. Images were collected from 4-6 sections/mouse (minimum 20 images/mouse). Quantification was performed on background subtracted images using ImageJ. Automatic thresholding was applied to GAD65/67 images before using the puncta analyzer function on NeuN-outlined neuronal cell bodies.
